# Consortium bibliometric analysis of hotspots and frontiers of immunotherapy in pancreatic cancer

**DOI:** 10.1101/2022.11.10.515943

**Authors:** Qiong Xu, Yan Zhou, Heng Zhang, Haipeng Li, Haoren Qin, Hui Wang

## Abstract

**Background:** Pancreatic cancer is one of the most common malignant neoplasms with an increasing incidence, low rate of early diagnosis, and high degree of malignancy. In recent years, immunotherapy has made remarkable achievements in various cancer types including pancreatic cancer, due to the long-lasting antitumor responses elicited in human body. Immunotherapy mainly relies on mobilizing the host’s natural defense mechanisms to regulate the body state and exert anti-tumor effects. However, no bibliometric research about pancreatic cancer immunotherapy has been reported to date. This study aimed to assess research trends and offer possible new research directions in pancreatic cancer immunotherapy.

**Methods:** The articles and reviews related to the pancreatic cancer immunotherapy were collected from the Web of Science Core Collection. CiteSpace, VOSviewer and an online platform were used to analyse co-authorship, citation, co-citation, and co-occurrence of terms retrieved from literatures highlighting the scientific advances in pancreatic cancer immunotherapy.

**Results:** We collected 2475 publications and the number of articles was growing year by year. The United States had a strong presence worldwide with the most articles. The most contributing institution was Johns Hopkins University (103 papers). EM Jaffee was the most productive researcher with 43 papers, L Zheng and RH Vonderheide ranked second and third, with 34 and 29 papers, respectively. All the keywords were grouped into four clusters: “immunotherapy”, “clinical treatment study”, “tumor immune cell expression”, “tumor microenvironment”. In the light of promising hotspots, keywords with recent citation bursts can be summarized into four aspects: immune microenvironment, adaptive immunotherapy, immunotherapy combinations, and molecular and gene therapy.

**Conclusions:** In recent decades, immunotherapy showed great promise for many cancer types, so various immunotherapy approaches have been introduced to treat pancreatic cancer. Understanding the mechanisms of immunosuppressive microenvironment, eliminating immune suppression and blocking immune checkpoints, and combining traditional treatments will be hotspots for future research.

## Introduction

Pancreatic cancer is a common malignant tumor of digestive system and is one of the leading causes of cancer-related deaths today, which is principally seen in men and in advanced age groups (40-85 years) [1]. Although there are many causes for pancreatic cancer, smoking and family history appear to be dominant factors [2]. Despite advances in surgery, chemotherapy and radiotherapy, mortality rate remains high, and 5-year survival rate do not exceed 20%-25% even after surgical resection. Many patients are already in advanced stages at the time of diagnosis [2,3]. In this case, the focus of treatment has shifted to improve overall survival using immunotherapy, targeted therapy and endocrine therapy. Among those strategies, immunotherapy is one of the most promising.

Immunotherapy can boost natural defense to eliminate malignant tumor cells and it is a monumental breakthrough for cancer treatment [4]. Several clinical studies have shown immunotherapy works continuously on tumor cells [5-8]. The major categories of immunotherapy include immune checkpoint inhibitors (ICIs), cancer vaccines, cytokine therapies, adoptive cell transfer, and oncolytic virus therapies. ICIs have been applied in the treatments of many cancer types, including melanoma, lung cancer and renal-cell cancer [9]. In healthy individuals, the immune checkpoint regulates the actions of ligands and receptors to keep the immune functions of T cells normal and balanced. When T cells are activated, they will express more immune checkpoint receptors, such as programmed cell death protein 1 (PD-1) or cytotoxic T lymphocyte-associated antigen 4 (CTLA-4) [10,11]. In 2007, programmed cell death-ligand 1 (PD-L1) was first identified as a new pancreatic cancer prognostic indicator due to the confirmation of up-regulation of PD-L1 expression in human pancreatic cancer cell [12]. Anti-CTLA-4 and anti-PD-1/anti-PD-L1 agents can inhibit immune checkpoints, and make T cells activate and provide effective approaches for tumor immunotherapy [13]. However, several immunotherapies that have been successful in various types of cancer, failed to act on pancreatic cancer due to its low tumor mutation burden and immunosuppressive tumor microenvironment [14,15]. Many researches indicated that pancreatic cancer may evade the immune surveillance by inducing the development of immune-suppressive T cells, such as regulatory T (Treg) cells. Treg cells exert immunosuppressive functions through multiple mechanisms [16]. One mouse experiment of pancreatic cancer confirmed that the combination PD-1 inhibitors and OX40 agonist reduced the proportion of Treg and exhausted T cells, and increased the number of CD4+ and CD8+ T cells in pancreatic tumors [17]. Thus, an increasing amount of research on immunotherapy has been conducted in recent years, and the research tends to suggest the need to overcome immune suppression and combine immunotherapy with traditional strategies to treat pancreatic cancer in the future.

By revealing the structure of knowledge in a scientific field over a certain period of time, bibliometrics can identify research hotspots and explore cutting-edge trends in recent years [18,19]. However, the global bibliometric analysis on the knowledge mapping of immunotherapy for pancreatic cancer has not yet been performed.

This study is based on immune-related studies of pancreatic cancer included in the Web of Science Core Collection, and we used visualization analysis software CiteSpace and VOSviewer to generate scientific knowledge map. Through co-occurrence and cluster analysis of related literature research institutions, keywords, authors, among others, we explored the development hotspots and frontier trends of pancreatic cancer immunology research, and provide support and reference for the subsequent related research.

## Materials and methods

### Dataset selection

The Science Citation Index Expanded of Clarivate Analytics’ Web of Science Core Collection (WoSCC) was used as data source. The WoSCC database is regarded as one of the most comprehensive, systematic, and authoritative databases, which contains over 15000 influential high-quality journals throughout the world. According to previous studies, papers in the Web of Science Core Collection (WOSCC) can represent the status of medical science; therefore, we chose WOSCC as our data source. WoSCC has been widely used for bibliometrics analysis and visualization of scientific literature in a substantial number of studies [20-23]. We selected articles about pancreatic cancer immunotherapy on March 10, 2022. The main search terms were as follows: #1, “Immunotherap*” OR “Anti-CTLA-4” OR “Anti-PD-1” OR “Anti-PD-L1” OR “Ipilimumab” OR “Tremelimumab” OR “Pembrolizumab” OR “Atezolizumab” OR “GVAX” OR “Algenpantucel-L” OR “Dendritic cell vaccine” OR “K-Ras vaccine” OR “CAR T cell therapy” OR “GV1001 vaccine”; #2, “Pancreatic cancer*” OR “Pancreatic carcinoma” OR “Carcinoma of pancreas” OR “Cancer of pancreas” OR “Pancreas neoplasm*” OR “Pancreas cancer*”; #3, “#1” AND “#2”. The publications from January 2007 to December 2021 were searched. The language was limited to English, and the literature types were limited to “articles” and “review” articles.

### Data visualization and analysis

Data were downloaded and analyzed independently by two researchers to ensure its accuracy and reliability. When the two researchers had different opinions regarding the data, a third researcher decided the prevailing opinion. The data analysis process strictly followed the corresponding statistical process. We used VOS viewer (version 1.6.18) [19], Citespace (version 5.8.R3) [18], and an online platform for bibliometric analysis (http://bibliometric.com) to construct visualization networks between researchers, journals, institutions and countries. Co-authorship, citation, co-citation, and co-occurrence analysis were also achieved. In VOSviewer, nodes represent different objects, such as countries/regions, institutions and researchers. The circle size of nodes represented frequency. Meanwhile, CiteSpace was used for the co-citation analysis of references and the extraction of keywords and references with high citation bursts. Also, CiteSpace generated a dual-map overlay of journals. The setting parameters were as follows: top 30 for selection criteria, minimum spanning tree and pruning sliced networks for pruning, and one or two years per slice from 2007 to 2021. In addition, the visualization of global publications and citation trends was analyzed by the online citation report of WoS.

## Results

### Trend of Global Publications and Citations

In the first place, 2812 papers were searched out. After excluding non-English and irrelevant articles, a total of 2475 documents (1702 articles and 773 reviews) were included in the study (Fig 1).

**Fig 1.**
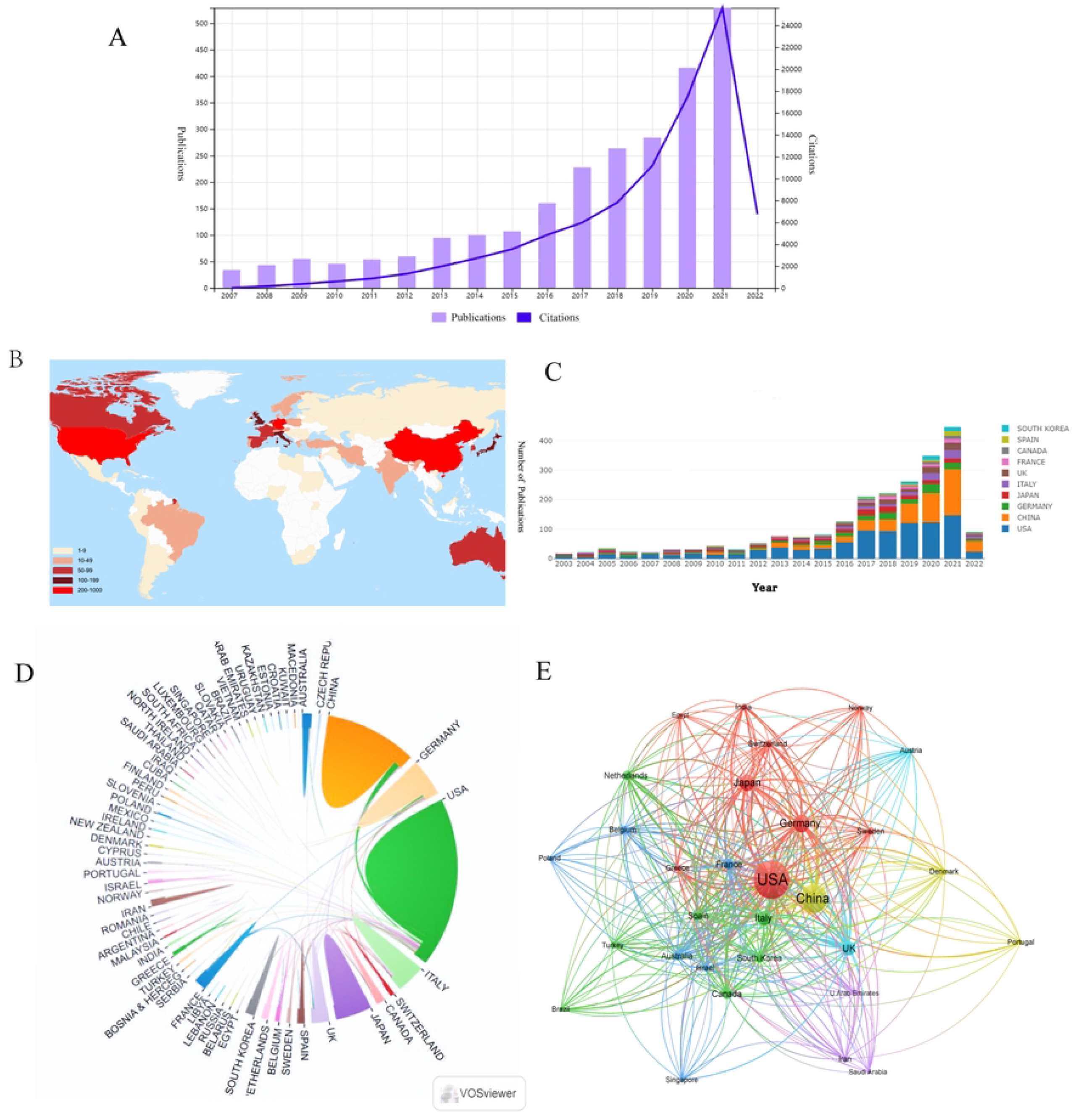
Schematic diagram of the search process.

According to Fig 2A, we see that the number of papers gradually increased from 34 in 2007 to 529 in 2021. From 2014, more than 100 articles were published every year. In the last five years, this area has developed rapidly and published 1721 (69.54% of 2475) articles. There were 91208 citations for all papers, with an average of 36.85 citations per paper.

**Fig 2.**
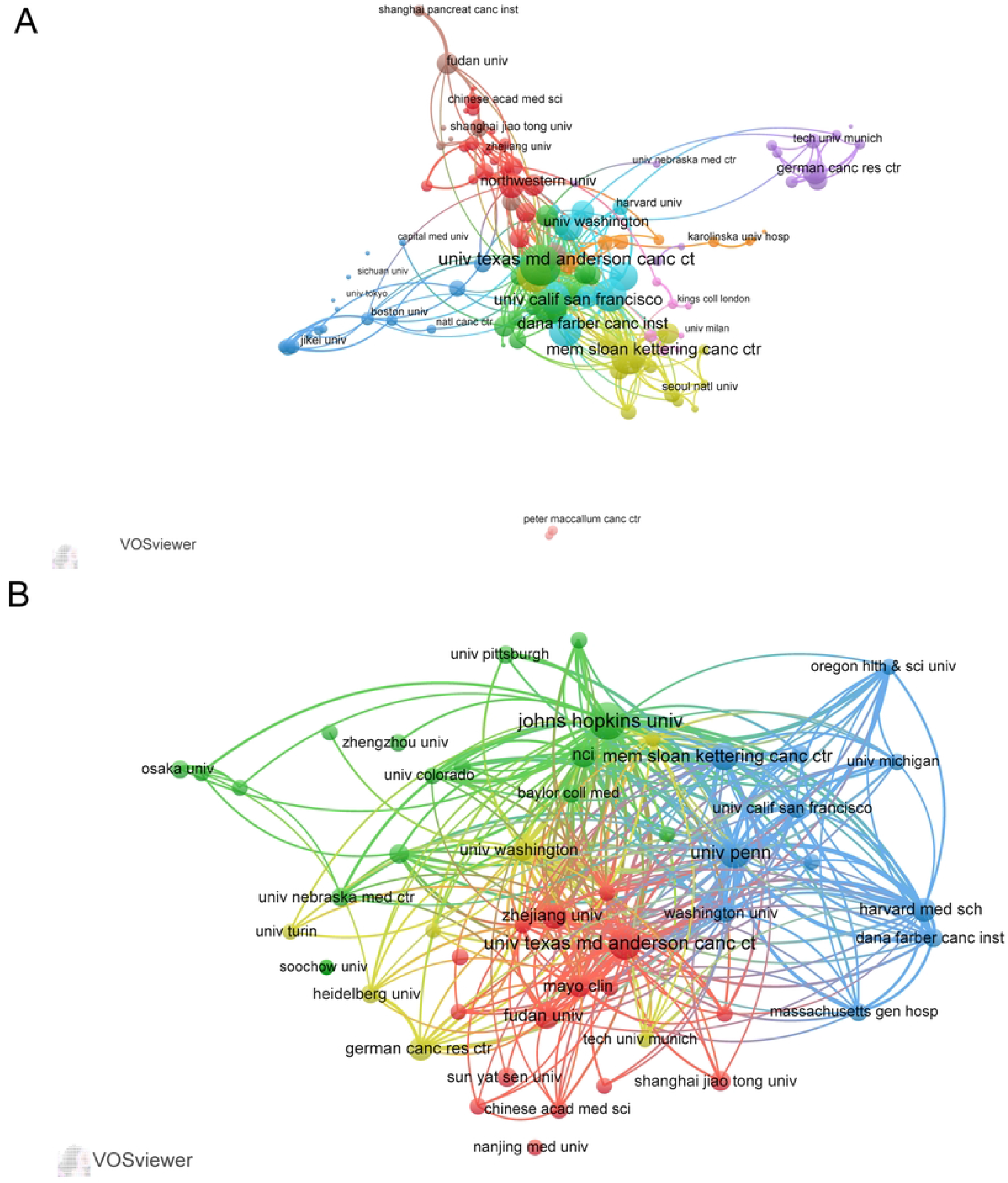
A: The annual number and total citations of articles related to pancreatic cancer immunotherapy published from 2007 to 2021. B: World map based on the number of articles of countries/regions. C: The trend of annual number of articles from top ten countries/regions. D: The international collaborations between countries/regions. The links between countries/regions represented the frequency of the collaborations. E: The citation network between top 30 countries/regions in terms of articles quantity, generated by VOSviewer. The size of nodes indicated the number of articles, while the thickness of links indicated the citation strength.

### Country/Region and Institution Analyses

A total of 72 countries/regions had published articles on pancreatic cancer immunotherapy (Fig 2B). Figure 2C shows the trend in the number of articles from the top 10 countries. The United States (US) was the most productive country, with 1096 publications (Table 1), followed by China (567), Germany (222). Additionally, the H-index of the US far exceeded that of China. Although China was ranked as the second most productive country, the average number of citations per article (18.63) was lower than those for Germany (37.35), Japan (33.97), and Italy (43.33), indicating a relatively low impact of Chinese publications and a lack of high quality publications. Furthermore, the collaboration analysis according to country/region revealed that the US collaborated with many countries, including China and Italy. In contrast, collaboration among other countries was weak (Fig 2D). The citation network map in Fig 2E shows the citation relationships among the top 30 countries/regions in terms of the number of articles.

**Table 1.**
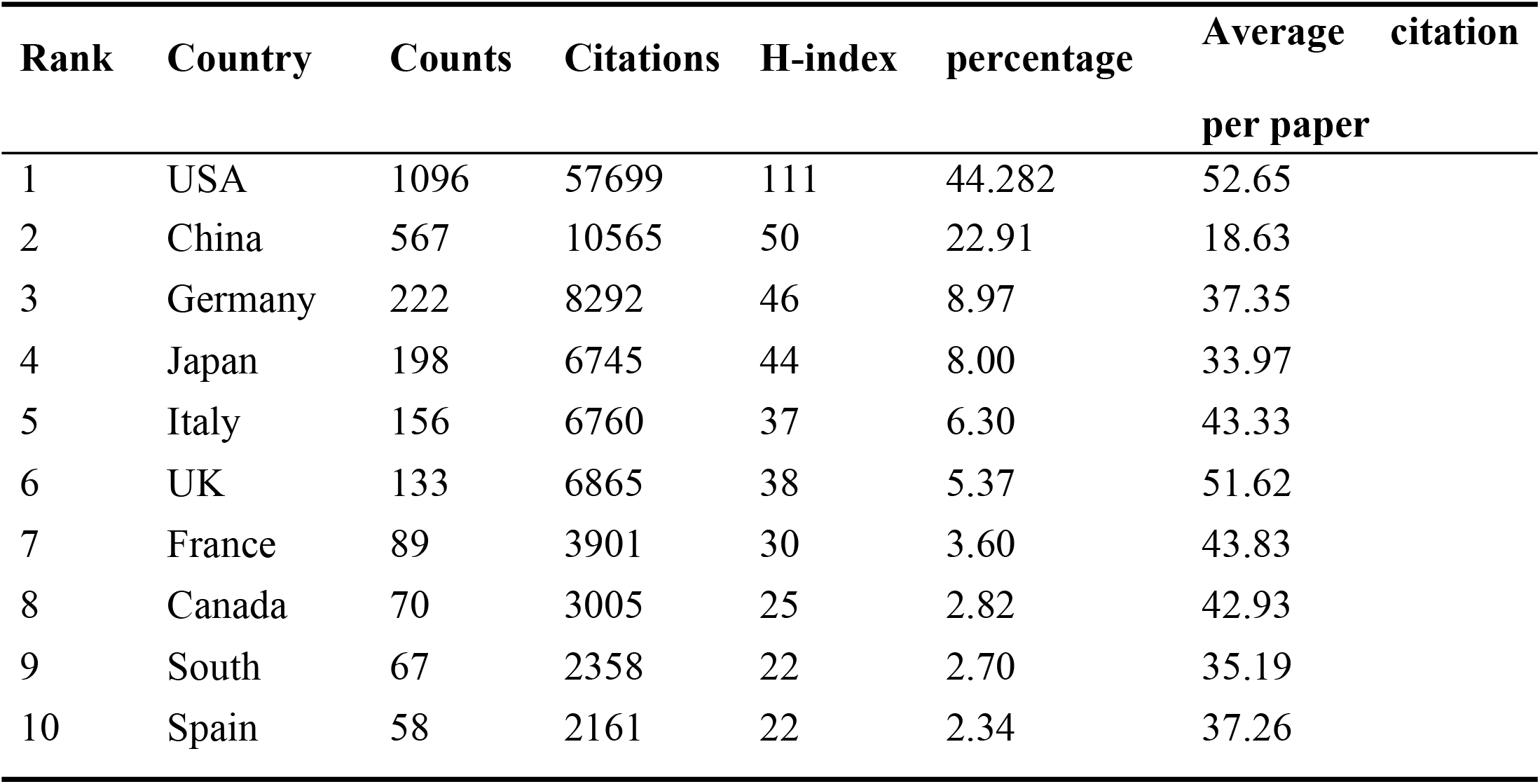
Top 10 productive countries/regions.

A total of 2880 institutions contributed to the field of pancreatic cancer immunotherapy, with the US accounting for seven out of 10 institutions (Table 2). Johns Hopkins University had the highest contribution, with 103 articles; the second-rank institution was the University of Texas MD Anderson Cancer Center (80 articles), followed by the University of Pennsylvania (70 articles). In the collaboration analysis according to institution (Fig 3A), we only selected institutions with 10 or more articles; accordingly, 141 institutions were analyzed. The collaboration analysis organized these institutions into 10 clusters, with especially close collaboration within each cluster. The institution citation analysis revealed that the top three institutions in terms of the total link strength (TLS) were the University of Pennsylvania (TLS=2384), Johns Hopkins University (TLS=2313), and University of Texas MD Anderson Cancer Center (TLS =1533) (Fig 3B, Table 2).

**Table 2.**
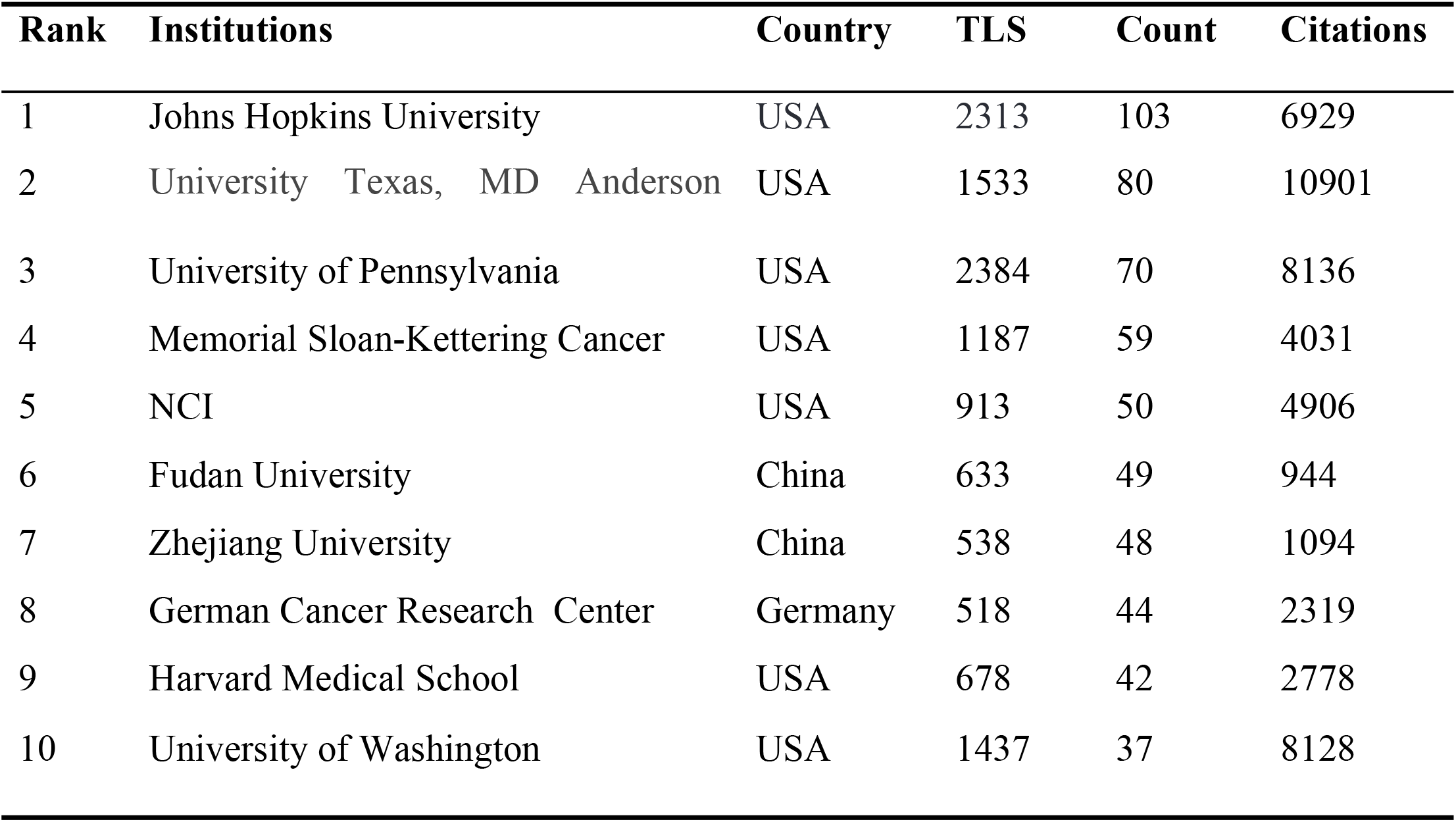
Top 10 productive institutes.

| Rank | Institutions | Country | TLS | Count | Citations |
| --- | --- | --- | --- | --- | --- |
| 1 | Johns Hopkins University | USA | 2313 | 103 | 6929 |
| 2 | University Texas, MD Anderson | USA | 1533 | 80 | 10901 |
| 3 | University of Pennsylvania | USA | 2384 | 70 | 8136 |
| 4 | Memorial Sloan-Kettering Cancer | USA | 1187 | 59 | 4031 |
| 5 | NCI | USA | 913 | 50 | 4906 |
| 6 | Fudan University | China | 633 | 49 | 944 |
| 7 | Zhejiang University | China | 538 | 48 | 1094 |
| 8 | German Cancer Research Center | Germany | 518 | 44 | 2319 |
| 9 | Harvard Medical School | USA | 678 | 42 | 2778 |
| 10 | University of Washington | USA | 1437 | 37 | 8128 |

**Fig 3.**
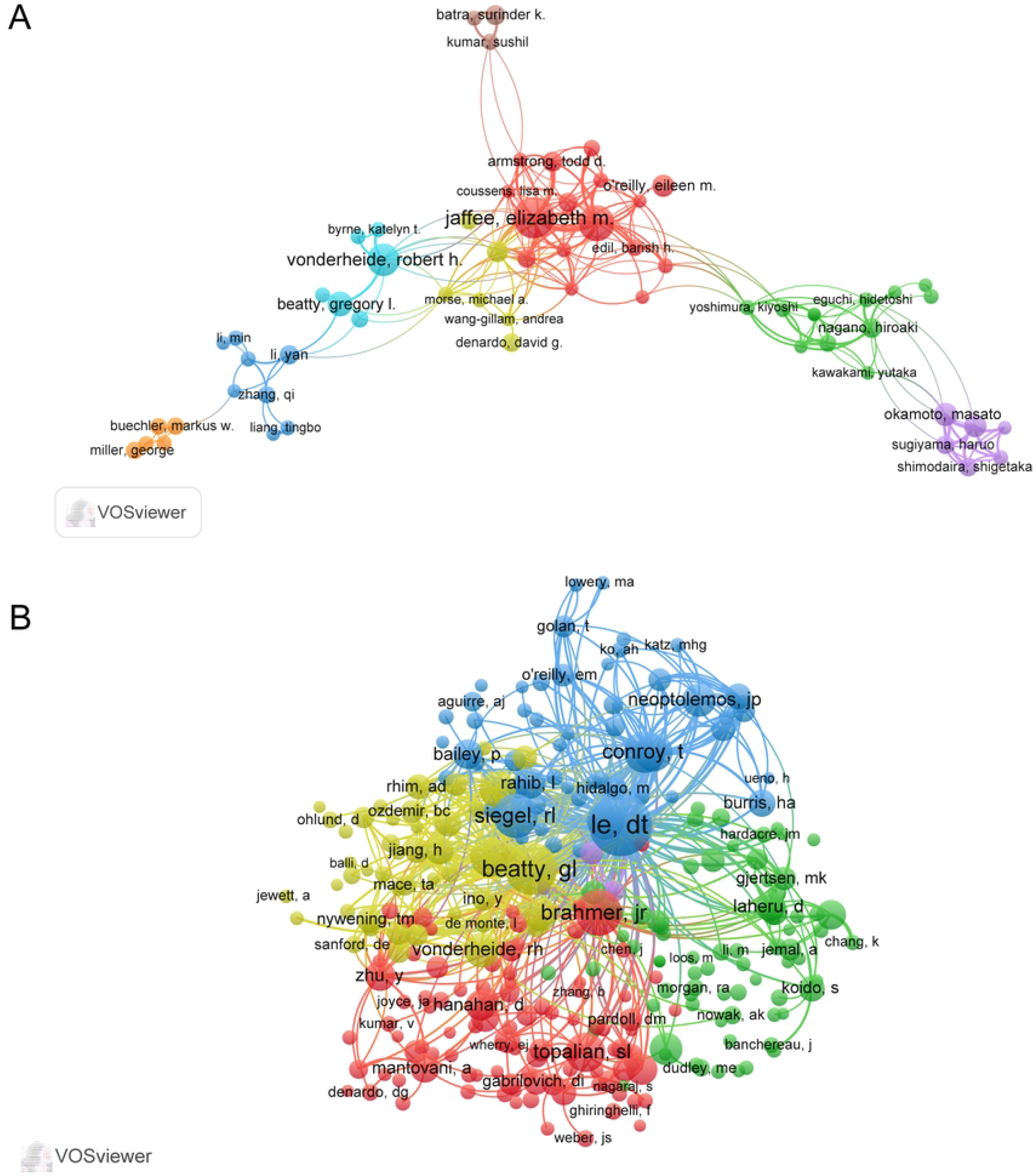
A: The cooperation map of institutions performed with VOS viewer. The size of nodes represented the number of publications, while the thickness of the lines indicated the collaboration strength. B: The citation network of institutions created with VOS viewer.

### Authors and Co-cited Authors

In total, 14998 authors and 59839 co-cited authors were identified in this study. Table 3 shows the 10 most prolific authors and the top ten co-cited authors with the largest TLS. EM Jaffee was the most productive author, with 43 articles in the field of pancreatic cancer immunotherapy, followed by L Zheng and RH Vonderheide, with 34 and 29 articles, respectively. The collaboration analysis revealed that authors generally did not collaborate closely (Fig 4A). EM Jaffee and L Zheng were placed at central positions in the clusters, and collaborated closely with each other. However, L Zheng and RH Vonderheide lacked collaboration. In the co-citation analysis, there were 5 clusters, 277 nodes, and 33190 links in the network map (Fig 4B). DT Le had the highest centrality, and had close contact with other scholars. The top three authors in terms of the TLS were DT Le (TLS=20853), GL Beatty (TLS=17327), and JR Brahmer (TLS=10346).

**Table 3.** The top ten authors and co-cited authors.

| Rank | Author | Docume | Citations | Co-cited | Citations | TLS |
| --- | --- | --- | --- | --- | --- | --- |
| 1 | Jaffee, E. M. | 43 | 3598 | Le, D.T. | 749 | 20853 |
| 2 | Zheng, L | 34 | 2095 | Beatty, G. L. | 550 | 17327 |
| 3 | Vonderheide, R. H. | 29 | 5573 | Brahmer, J.R. | 451 | 10346 |
| 4 | Beatty, G. L. | 18 | 2487 | Siegel, R.L. | 414 | 6779 |
| 5 | Novelli, F | 18 | 709 | Conroy, T | 379 | 9157 |
| 6 | Cappello, P | 17 | 683 | Topalian, S.L. | 347 | 6582 |
| 7 | Okamoto, M | 17 | 349 | Feig, C | 339 | 9204 |
| 8 | Koido, S | 16 | 348 | Royal, R.E. | 324 | 8797 |
| 9 | Oreilly, E.M. | 15 | 544 | Vonhoff, D.D. | 324 | 7848 |
| 10 | Le, D.T. | 14 | 2150 | Hingorani, | 301 | 7750 |

**Figure 4.**
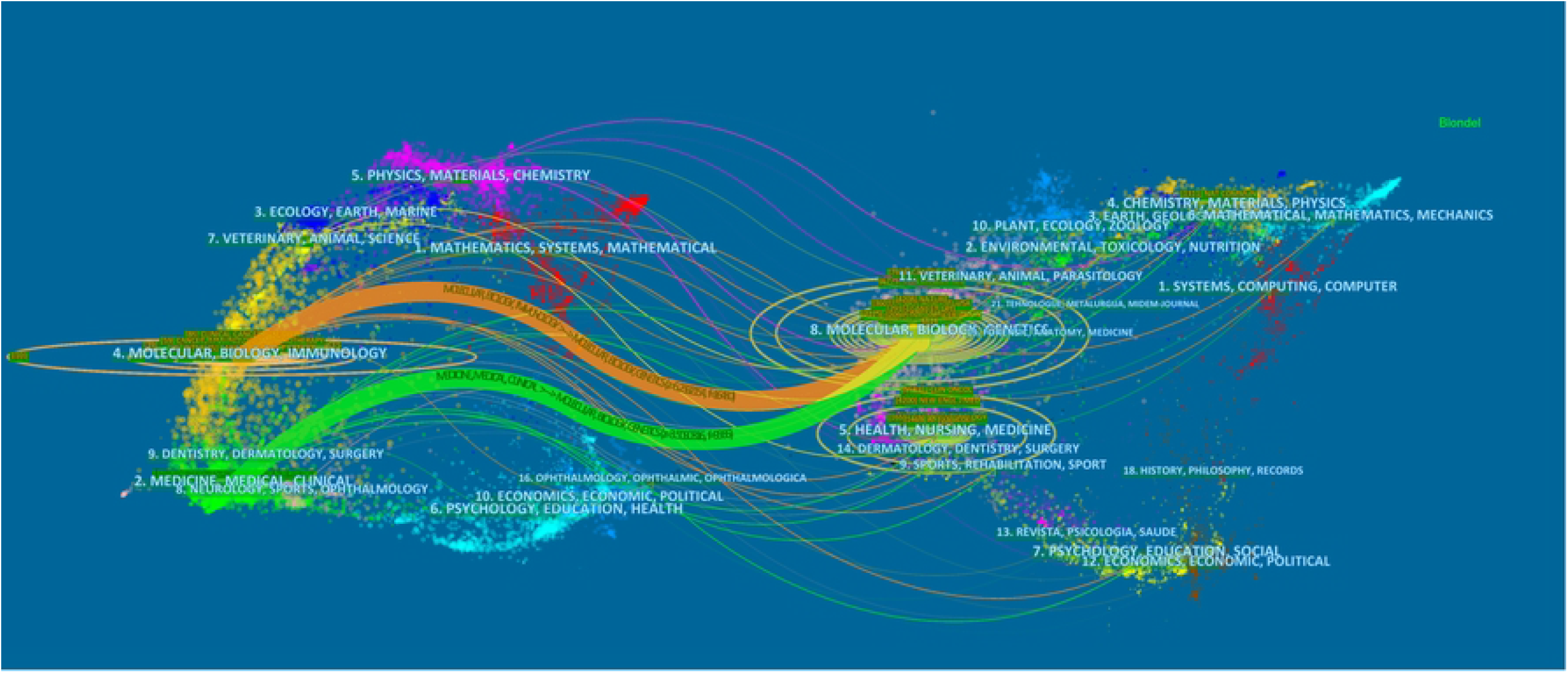
The cooperation map of authors (A) and co-citation map of authors (B) conducted with VOSviewer.

### Contributions of Top Journals

A total of 612 academic journals published articles in the field of pancreatic cancer immunotherapy from 2007 to 2021, including 111 journals with more than 5 articles. Cancers was the most active journal, with 99 articles in the field; however, their impact factor (IF) was much lower than that of the second-ranked journal, Clinical Cancer Research (6.639 vs 12.53). The journal with the third highest number of publications was Oncoimmunology (69 articles, IF=8.11). The Journal for Immunotherapy of cancer had the largest IF (13.75), followed by Clinical Cancer Research (12.53) and Cancer Research (12.701). The majority of the top ten journals were classified as Q1 or Q2 according to Journal Citation Report 2020 criteria. Half of the top 10 journals derived from the US.

Fig 5 shows a dual-map overlay of academic journals, with citation relationships between citing journals (left half of the map) and cited journals (right half of the map). In Fig 5 and Table 4, the labels indicate the fields covered by the journal, the colored lines depict different citation paths, and the path width is proportional to the z-score scale. We noted two main citation paths, the orange path and the green path, which indicated that articles from molecular, biology and genetics journals were frequently cited by articles from of molecular, biology, immunology, medicine, medical, and clinical journals.

**Table 4.**
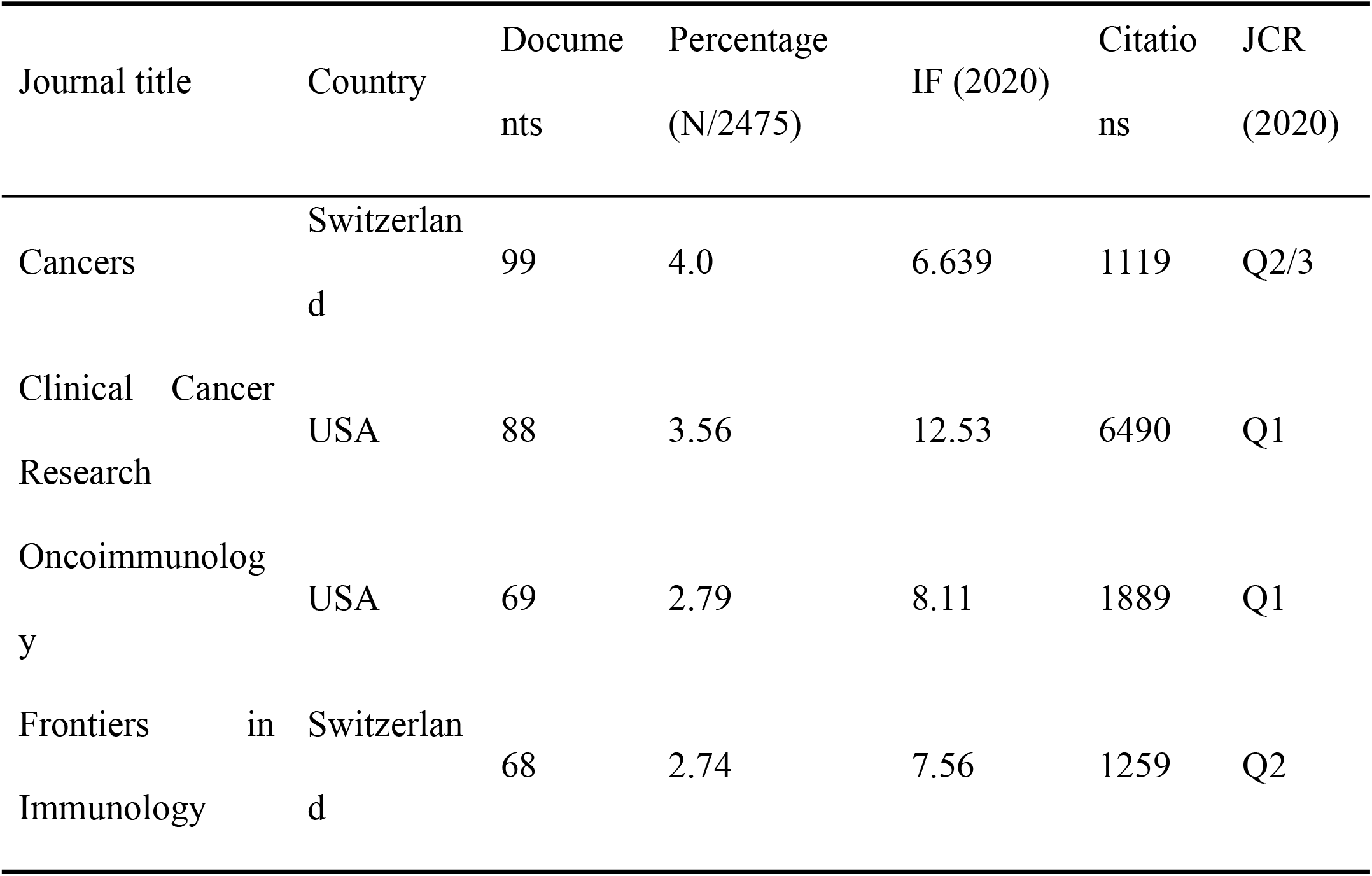
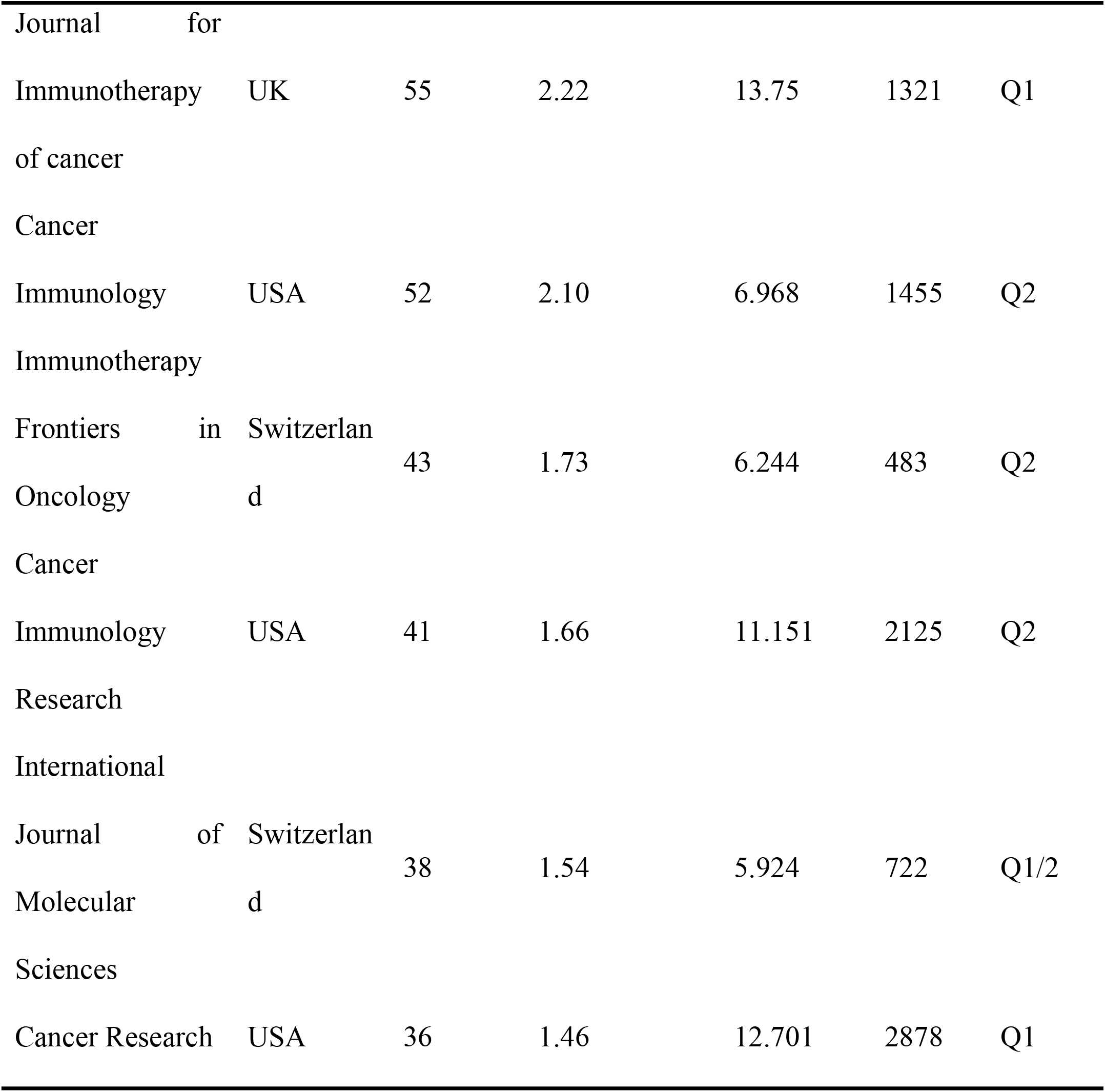
The top ten most active journals.

**Figure 5.**
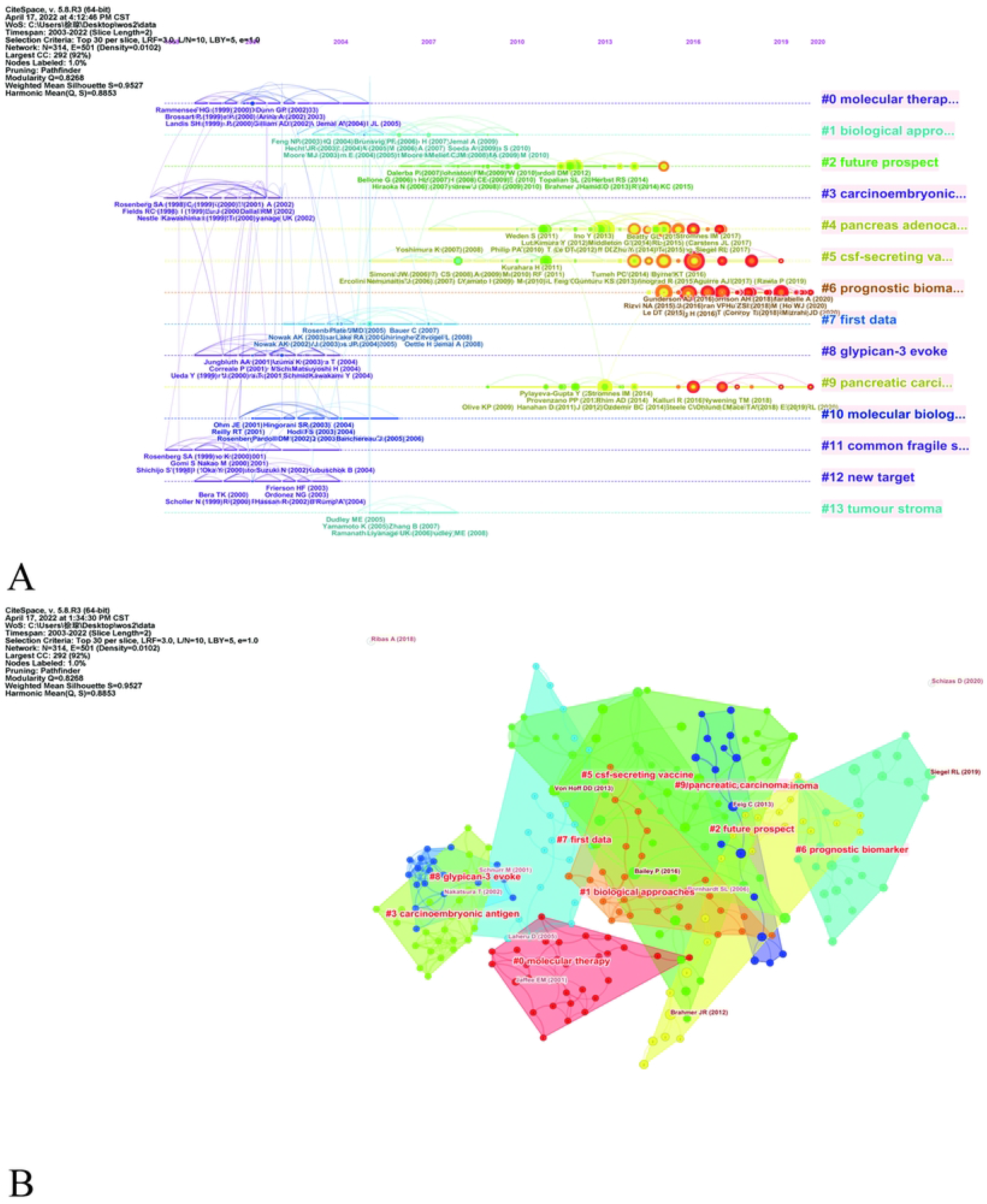
A dual-map overlap of journals on immunotherapy for pancreatic cancer completed by CiteSpace.

### Analysis of Co-cited References

Table 5 shows the top ten co-cited references on pancreatic cancer immunotherapy. “Safety and activity of anti-PD-L1 antibody in patients with advanced cancer” by Brahmer et al. [9] was the highest cited article with 387 citations. This article mainly found that antibody-mediated PD-L1 blockade induced durable tumor regression and long-term disease stabilization in patients with advanced cancers. The second and third articles were by Royal et al. [24] and Conroy et al. [25] and had 319 and 291 citations, respectively. In the co-citation network analysis of the references, the references were divided into 10 clusters, the weighted mean silhouette and modularity Q were both greater than 0.8, and the clustering structure was reasonable (Fig 6A). The top three clusters were “#0 molecular therapy”, “#1 biological approaches”, and “#2 future prospect”.

**Table 5.**
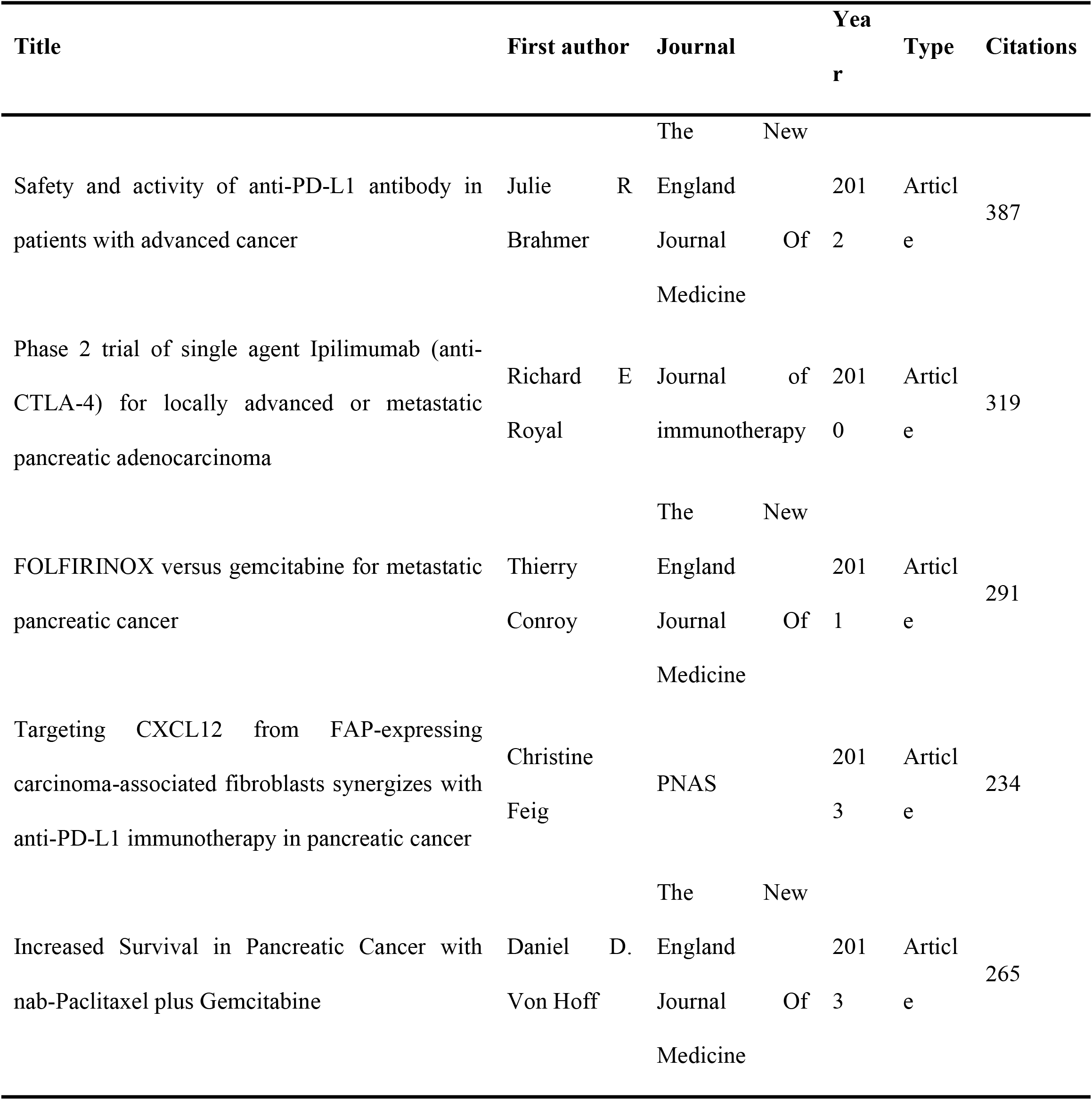
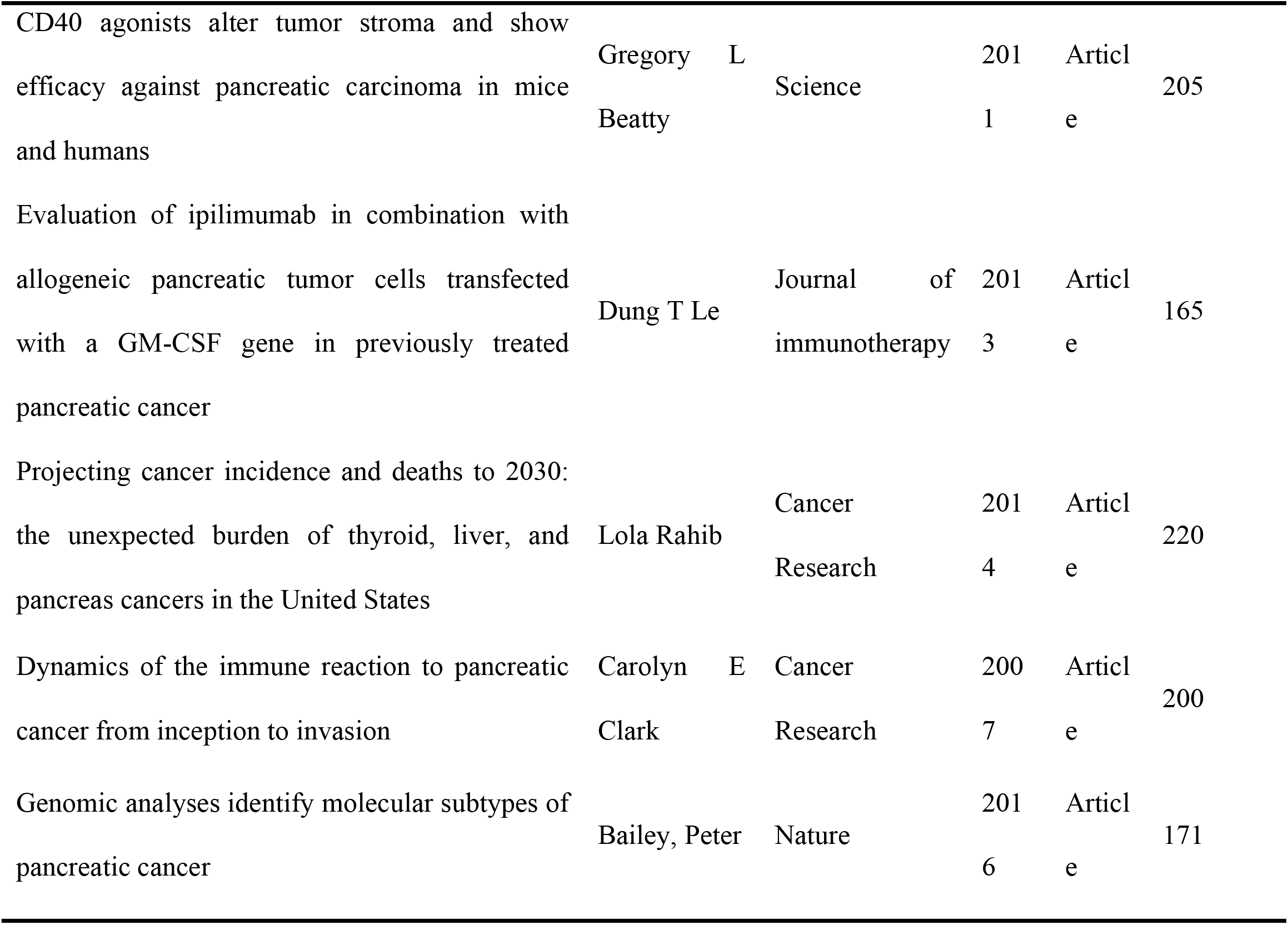
Top 10 co-cited references concerning the research.

**Fig 6.**
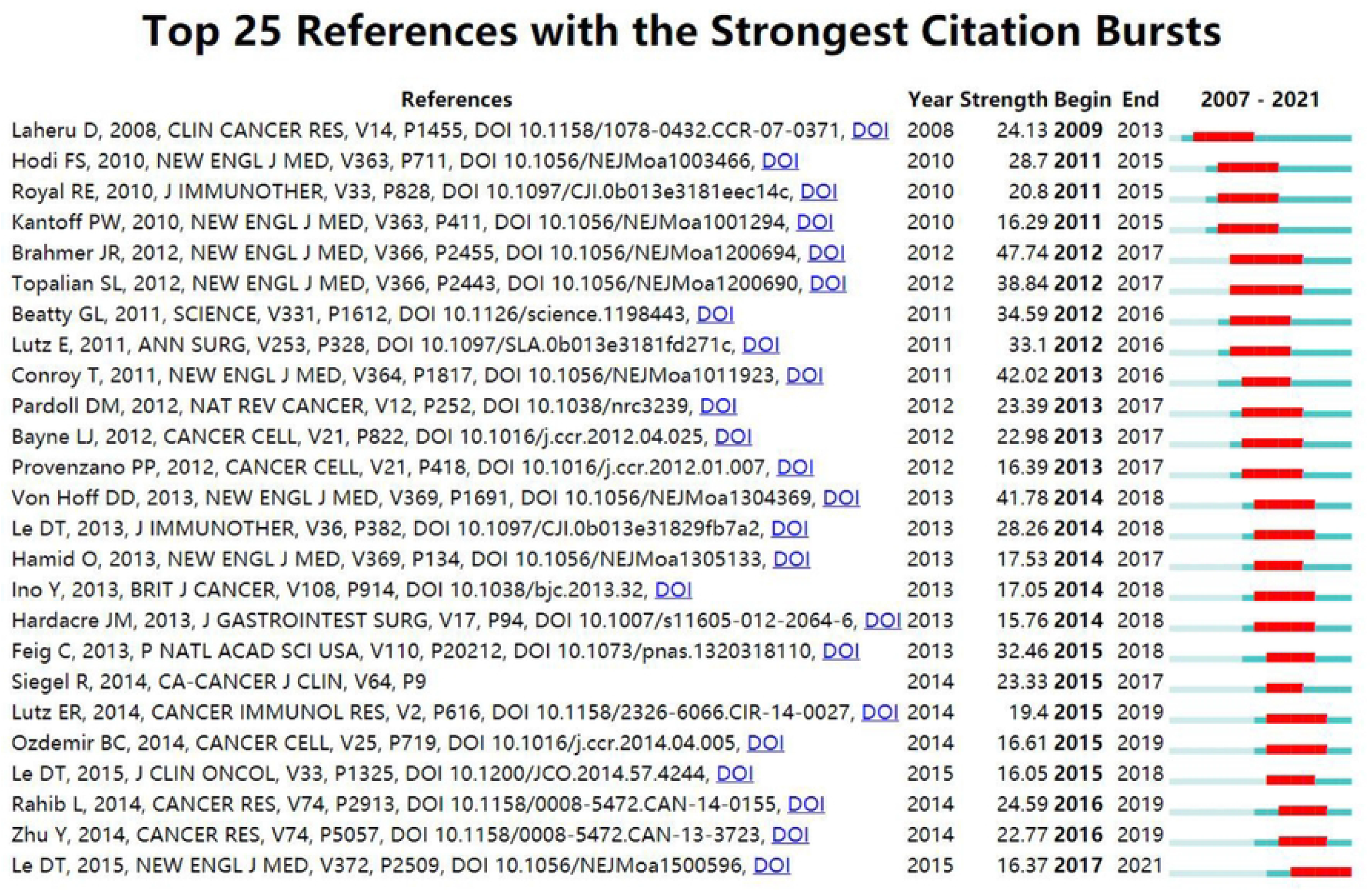
Timeline view (A) and Cluster view (B) of co-citation references related to pancreatic cancer immunotherapy visualized by CiteSpace. Fifteen labeled clusters are colored on the right. The nodes on the line represented the cited references.

We also performed a timeline analysis of co-cited references to reveal the changes in research hotspots over time (Fig 6B). The vertical axis in Fig 6B represents the clustering label, with a total of 13 clusters, and the horizontal axis shows the timing of the occurrence of important reference nodes. “#3 carcinoembryonic antigen” and “#11 common fragile site” were the earliest hotspots, and there were relatively few studies on molecular therapy in recent years. The most recent research hotspots were “#6 prognostic biomarker” and “#5 csf-secreting vaccine”, suggesting that concerns are shifting more toward cancer genes and immunotherapy research.

“Citation bursts” refer to references well known in the field and widely cited during a particular period. CiteSpace identified 15 most frequently cited references (Fig 7). In Fig 7, the red segmented lines indicate the active period. A short citation burst in the field of pancreatic cancer immunotherapy began in 2011, corresponding to the advent of immunotherapy for pancreatic cancer. Other bursts emerged later, and have developed relatively quickly in recent years. The most recent highly cited references (2019-2021) were Rahib et al. [26], Zhu et al. [27], and Le et al. [28], and these bursts are still ongoing.

**Fig 7.**
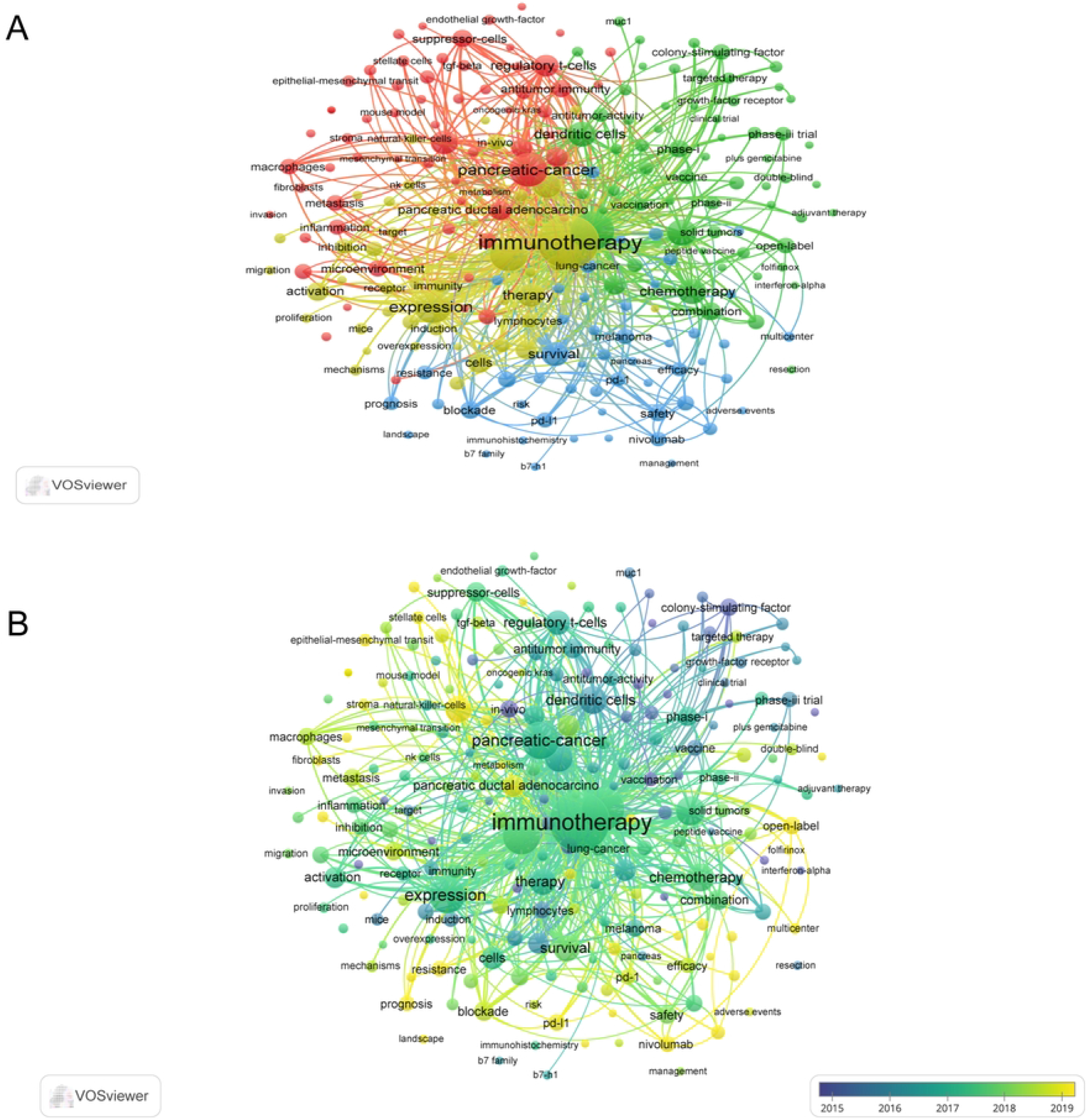
The top 25 references with the strongest citation bursts during 2007 to 2021.

### Keywords Analysis of Research Hotspots

We used VOSviewer to create a keyword network map, overlay map, and density map. There were a total of 7815 keywords in 2475 articles, of which, 223 keywords occurred 20 or more times and 30 keywords occurred more than 100 times (Fig 9B). Keywords that appeared at least 20 times were analyzed. Immunotherapy and pancreatic cancer were the most frequently occurring keywords, consistent with the objectives of this study (Fig 9B).

In the keyword co-occurrence analysis, We found that the last 15 years of pancreatic cancer immunotherapy research can be summed up in the following four main topics that had the highest number of publications: “immunotherapy”, “clinical treatment study”, “tumor immune cell expression”, “tumor microenvironment”. In addition, each topic covered many small branches, as evident from the colors of each main topic indicated by large circles and related smaller circles of the same color. (Fig 8A). In Figure 8A, the main blue-marked cluster “immunotherapy” can be subdivided into the branches “t-cell immunity”, “PD-L1”, “nivolumab”, “pembrolizumab”. The green cluster is related to “clinical treatment”, focusing on the clinical efficacy of conventional treatment, with keywords such as “gemcitabine”, “radiation therapy”, “adjuvant therapy”, and “targeted therapy”. The yellow cluster is mainly related to anti-tumor effects of immune cells and biological effect of immunotherapy, which may aid in elucidating the mechanisms of pancreatic cancer immunity. The red cluster is related to the “tumor microenvironment”, with keywords such as “tumor-associated macrophage”, “suppressor T cell”, “muc-1”, and “regulatory T-cell”.

**Fig 8.**
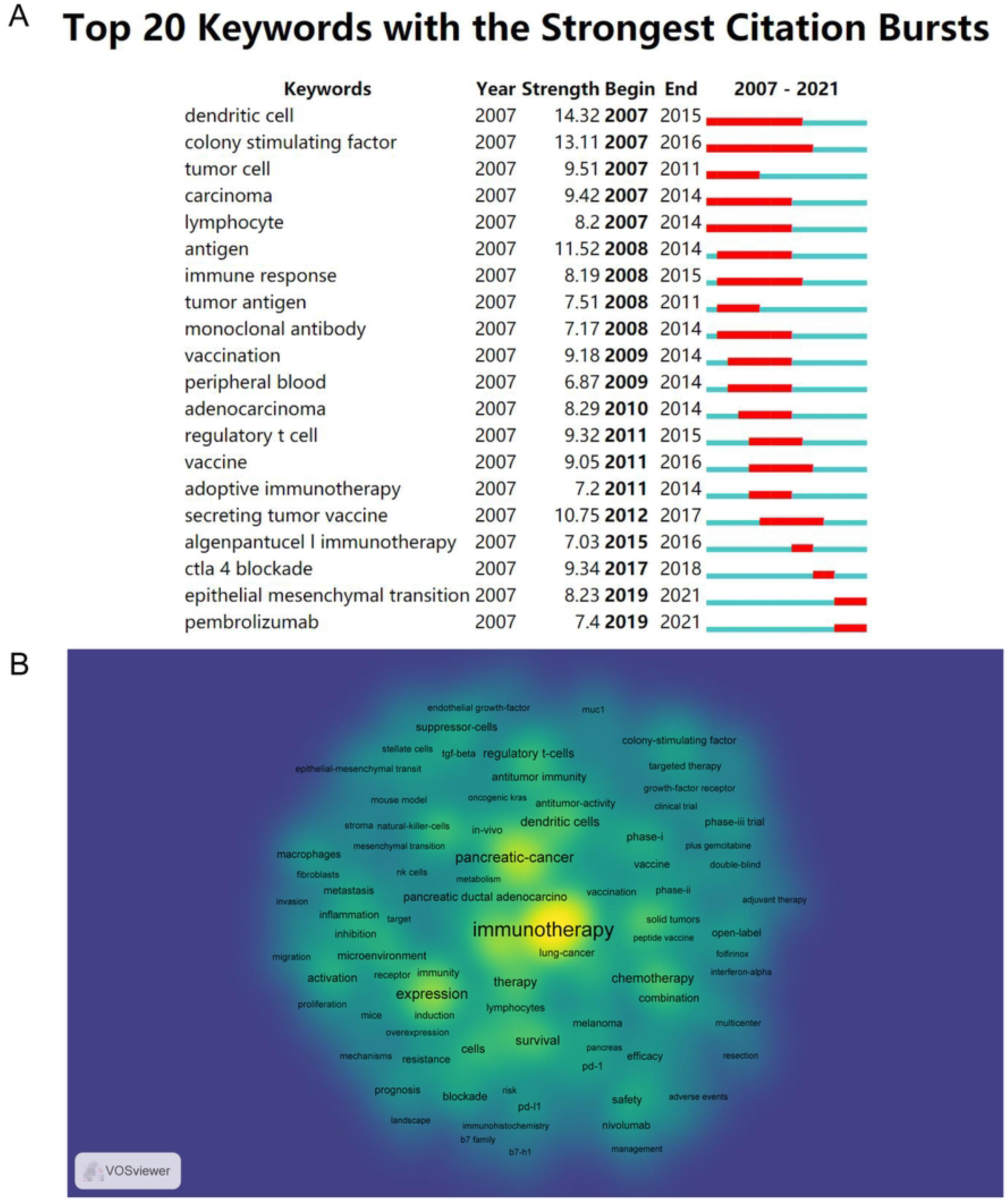
(A) The network of 223 keywords with a frequency of no fewer than 20 times. All keywords could be grouped into four clusters. (B) The overlay visualization map showed that keywords are colored according to their average occurrence time.

In Fig 8B, keywords are colored according to the average year of their occurrence in an overlay map. In the past decade, “nivolumab”, “vaccine”, and “gemcitabine” appeared frequently, suggesting that immune clinical trials were initiated and traditional anti-cancer therapies were well researched. The tumor microenvironment and immune checkpoint inhibitors have been studied more recently, and may be the hotspots of future research.

Another method for identifying research hotspots is to focus on keywords with strong burst strength (Fig 9A). From 2007 to 2015, studies focused on anti-tumor immune mechanisms, as indicated by the high burst strength of keywords such as “dendritic cell”, “colony stimulating factor”, “lymphocyte”, “regulatory T cell”, etc. From 2015 to 2021, keywords such as “vaccine” and “immune checkpoint inhibitors” have high burst strength, suggesting that these areas are becoming new research hotspots.

**Fig 9.**
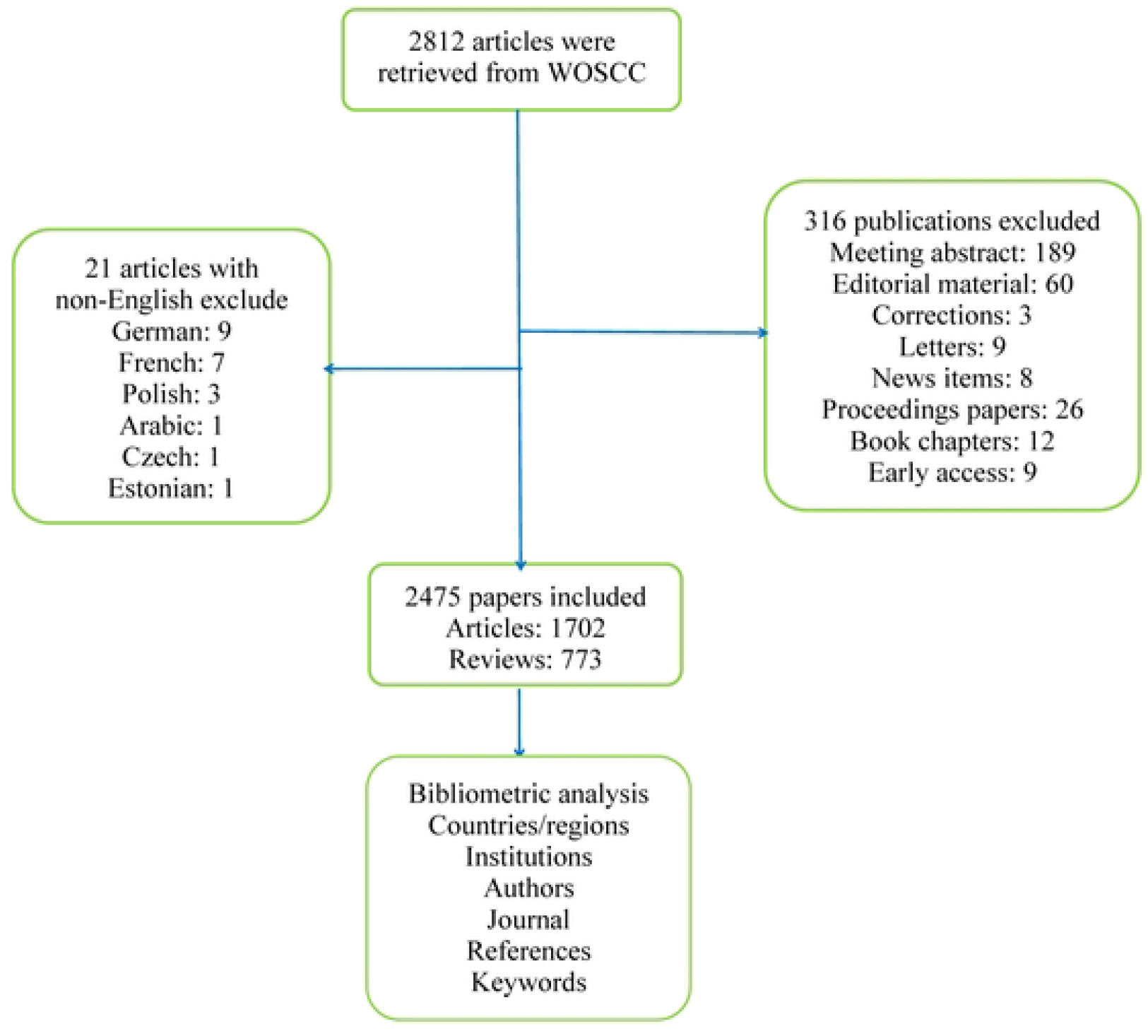
(A) Top 20 keywords with the strong citation bursts on immunotherapy for pancreatic cancer (B) Density map of keywords generated by the VOS viewer.

## Conclusions

We live in an age of information explosion; the speed of information generation is accelerating worldwide [29]. In the past 10 years, big data has become one of the most commonly used terms in healthcare, education, and industrial sectors [30,31]. Medical big data and medical data mining are multidisciplinary research fields, combining medicine and computer science, and have received wide attention, with the rapid development of information technology [30,32]. In this study, VOSviewer and CiteSpace software were used to analyze 15 years of research in the field of pancreatic cancer immunotherapy. We systematically reviewed the development of trends in the field by analyzing the core authors, highly productive countries and institutions, core journals, and keyword clusters.

Over the past 15 years, the literature on pancreatic cancer immunotherapy has been increasing (Fig 2A). Thus, it seems safe to consider this area as entering a golden age in the coming years. The US had the highest productivity. However, in recent years, there has been a gradual increase in the number of publications from China, Germany, and Japan, indicating a growing interest in immunotherapy for pancreatic cancer in these countries (Fig 2B and 2C). Thus, we boldly predict that more publications on immunotherapy for pancreatic cancer will appear in the future as a result of growing concern.

The H-index was proposed by Ali in 2005 to evaluate the quantity and quality of the academic output of researchers, and measure the impact of their scientific contributions [33]. “Total citations” refers to the number of times that an article has been cited in other publications (e.g. articles, reports, etc.). The present analysis found that the US ranked first in influence, (Table 1 and 2), thus, we concluded that the US is currently dominating this field. A late start in tumor immunology research may be the reason why China has lagged behind the US in this field. A co-citation analysis can depict the knowledge structure of the research area, identify research trends in the field, uncover cutting-edge research, and highlight high-impact discoveries [34]. DT Le was one of the most highly cited authors (Table 3), suggesting that this author wrote articles with greater academic impact. DT Le mainly focused on various immunotherapies, combining immunotherapy with other treatments, and exploring new treatments for tumors [35-38]. It should be noted that although China ranked second in terms of publications, only a few Chinese authors were highly cited by their peers.

The number of citations is a good indicator for evaluating an author’s academic influence [39]. It can be seen in Table 5 that the 10 most cited studies were published between 2007 and 2021, and the most cited article was by JR Brahmer; the most striking result of this article was the confirmation of a prominent role for anti-PD-L1 in many advanced tumors [40]. Table 6 shows a timeline view of co-cited references, which revealed developmental trends and changes in clustering across different periods. “#6 prognostic biomarker” is currently the most popular research topic.

In bibliometrics, keywords are used to understand how a field has developed. In Fig 8A, we noted that keywords related to pancreatic cancer immunotherapy were clustered into four groups. Fig 8B shows the time and frequency of the occurrence of keywords; early studies were focused on the immune system and its components (e.g. “regulatory T cell”, “dendritic cells”), and adaptive immunotherapy (e.g. “vaccine”). With the gradual deepening of immunization research, the immune microenvironment and clinical therapies are increasingly studied, and immunotherapy is attracting the attention of many researchers. We wish to ensure that more researchers pay attention to pancreatic cancer immunotherapy in the future, enabling new breakthroughs in the diagnosis and treatment of pancreatic cancer.

CiteSpace was used to find keywords with citation bursts, in an effort to reveal future trends in the field of pancreatic cancer immunotherapy. According to Fig 8B and 9A, the colors of all keywords were divided by VOSviewer according to the average publication year (APY). The latest keyword was “immune checkpoint inhibitors” (cluster 3, APY: 2019.91), followed by “mismath repair deficiency” (cluster 3, APY: 2019.84) and “tumor microenvironment” (cluster 1, APY: 2019.27). Besides, “biockade” (cluster 3, APY: 2018.54), and “tumor-associated macrophages” (cluster 1, APY: 2018.70) were the recent major topics in this field. Compared with Fig 8B and 9A, the study of immune checkpoint blockade was relatively the latest, suggesting that clinical and mechanistic studies of pancreatic cancer immunity are developing together.

### Research Hot spots

Furthermore, based on our analysis of keywords with the latest APY, “tumor microenvironment”, “adaptive immunotherapy”, “immunotherapy combinations”, and “molecular and gene therapy” may become research hotspots in the coming years.

### Tumor Microenvironment

Pancreatic cancer tumors are composed of malignant cells and extracellular matrix. Malignant cells only account for a small portion of tumor components, the rest is composed of fibroblasts, extracellular matrix, endothelial cells, and hematopoietic cells. Additionally, most of the immune microenvironment of pancreatic cancer is composed of these cells [41]. Thus, the structure, as a physical barrier, protects pancreatic cancer cells against an effective delivery of chemotherapeutic agents [42]. Carcinogenesis of pancreatic cancer involves progressive accumulation of driver mutations, including the oncogene K-Ras [43] and tumour suppressor gene TP53 [44]. K-Ras mutations are not only responsible for pancreatic cancer development, but also immediately track with other mutations, contributing to the aggressiveness of pancreatic tumors [45]. To have constant proliferation and survival, pancreatic cancer cells need continuous K-Ras signaling, and numerous downstream effector are engaged via K-Ras signaling. Accordingly, targeting K-Ras is likely to have a profound effect on pancreatic cancer. These data support the potential utility of targeting K-Ras or its downstream signaling pathways as a therapeutic approach in pancreatic cancer [46].

Morphological evolution of pancreatic cancer begins with the formation of precursor lesions, termed pancreatic intraepithelial neoplasia [47], with increasing histological grades followed by progression to invasive adenocarcinoma. As the cancer develops, it leads to changes in the surrounding tissue stroma. A key function of any non-transformed tissue stroma is to provide homeostatic response to injury with its immune, vascular and connective tissue components. However, cancer hijacks such physiological responses to create a favourable tumor microenvironment (TME) for its successful growth [48]. Over the past 10-15 years, numerous clinical and preclinical studies have provided abundant evidence that acellular and cellular components of the TME promote pancreatic tumorigenesis [49]. TME plays an important role in tumor development, metastasis, and resistance to chemotherapy [50], TME components can also result in the formation of connective tissue in primary and metastatic sites, or the promotion of the metastatic ability of pancreatic cancer by augmenting epithelial-mesenchymal transition (EMT) and angiogenesis [51]. The tumor microenvironment is receiving increasing attention as a target for tumor therapy [52]. Pancreatic cancer development is intertwined with multiple types of immunosuppressive cells, including regulatory T (Treg) cells, myeloid-derived suppressor cells (MDSCs) and tumour-associated macrophages (TAMs), and leads to an inherently immunosuppressed TME. Targeting immunosuppressive cells to modulate the immune TME is a current and future focal point of research interest.

### Adaptive Immunotherapy

In recent years, adaptive immunotherapy has dominated cancer immunology research through literature analysis [53]. The most popular of the adaptive immunotherapies are the immune checkpoint inhibitors, CAR-T, and cancer vaccines. Cancer vaccines include whole-cell vaccines, Dendritic Cell (DC), DNA and peptide vaccines that stimulate the presentation of immunogenic cancer antigens to the immune system, leading to activation of cancer antigen-specific CTLs in vivo [54] and subsequent anti-cancer immune response. Immune checkpoints are gaining attention as attractive targets for immunotherapy, Immune checkpoints are a regulatory mechanism used to modulate the T-cell immune response [55]. Anti-CTLA 4 and anti-PD-1/PD-L1 can block immune checkpoints and activate T cells, which is an important pathway for tumor immunotherapy [13]. Checkpoint inhibitors are most successful in tumor types with high PD-L1 expression and/or high microsatellite instability or mismatch repair deficiency [35,56]. Two FDA-approved monoclonal antibodies, pembrolizumab (KEYTRUDA) and nivolumab (OPDIVO), have been developed to block the interaction between PD-1 and its ligand. Two anti-PD-L1 antibodies, atezolizumab and avelumab, have also received FDA approval. Predictably, significant resources and efforts will be devoted to the continued development of adaptive immunotherapies in the future.

### Immunotherapy Combinations

Combination chemotherapy regimen consisting of oxaliplatin, irinotecan, fluorouracil, and leucovorin (FOLFIRINOX), as first-line therapy for patients with metastatic pancreatic cancer, showed a median overall survival of 11.1 months, which was greater than that with gemcitabine (6.8 months); however, the safety profile of FOLFIRINOX was less favorable than that of gemcitabine [25]. Chemotherapy alone has a low median survival and is more toxic. Additionally, the large number of immunosuppressive signals and effective immune evasion found in pancreatic cancer TME and low constitutive checkpoint expression, for patients with advanced pancreatic cancer, these drugs have minimal single agent activity [57] Many clinical studies have shown the ineffectiveness of immune checkpoint inhibitors alone [24,58]. Immunotherapeutic strategies that can overcome multiple barriers to effective antitumor immunity are needed; thus, the role of combination therapy is being explored in many trials and combination therapy has become a growing area of PDA research.

Pembrolizumab is a humanized monoclonal antibody that targets PD-1, thereby inhibiting the interactions of PD-1 with its ligands PD-L1 and PD-L2. Therefore, it is most often used in pancreatic cancer. Gemcitabine and paclitaxel, standard first-line chemotherapy regimens for metastatic pancreatic adenocarcinoma, have been previously reported to possess immunomodulatory properties [59]. A phase Ib/II study of gemcitabine, nab-paclitaxel, and pembrolizumab in metastatic pancreatic cancer (NCT02331251) reported that the median progression-free survival and overall survival of patients who completed treatment were 9.1 and 15.0 months, respectively, with a response rate of 71% (17 of 24 evaluable patients), including 2 complete responses [60]. Rational immunotherapeutic combinations may prove to be the optimal way to synergistically overcome the immunosuppressive TME and shift the balance to an anti-tumor TME.

The mechanical properties of EMT and subsets of pancreatic cancer with stem cells properties are increasingly recognized as effective modulators of therapeutic efficacy [61]. Pancreatic cancer stem cells have important functions during tumor recurrence and treatment resistance. EMT may promote tissue fibrosis and cancer metastasis [62,63]. A growing number of studies have clearly shown that cancer stem cells and EMT-type cells play an important role in chemical and radiological resistance [64-66]. Thus, this topic will likely become a trend and hotspot of future research.

### Molecular and Gene Therapy

According to the keyword cluster analysis, it can be seen that Molecular and gene therapy has become a research hotspot in recent years. Molecular profiling of cancer has identified potentially actionable drug targets, which has prompted attempts to discover clinically validated biomarkers to guide treatment decisions and enrollment in clinical trials [67]. Because all human cancers are primarily genetic diseases, identifying additional genes and signaling pathways could guide future research on this disease. The need to identify biomarkers that predict disease recurrence and druggable targets has driven our insights into the molecular structure of pancreatic cancer over the past 20 years of research. Despite many obstacles, the practice of guiding cancer treatment based on molecular aberrations is gaining momentum in oncology and has shown the potential to improve patient prognosis [51]. Jones S et al. examined the genetic makeup of human pancreatic cancer and identified KRAS, SMAD4, TP53, and CDKN2A as the four most frequently altered genes [68]. The research to date provides strong rationale for the integration of molecular profiling into clinical practice in the management of patients with pancreatic adenocarcinoma, to optimally plan a personalized therapeutic strategy through referral to specific studies of investigational targeted therapies or immunotherapies or to guide the optimal selection of standard cytotoxic chemotherapies [67].

## Strengths and Limitations

This study was the first to use a bibliometric approach to analyze global research trends in pancreatic cancer immunity over a 15-year period, which can aid scholars interested in pancreatic cancer immunity in establishing a clear framework of the existing research in the field and gaining insight into its development. Secondly, the clustering analysis of high-frequency keywords performed in this study can aid scholars in understanding the hotspots and focus of the field (tumor microenvironment, adoptive immunotherapy, etc.), and provide a reference for scholars in selecting topics.

However, this study also had some limitations. Firstly, bibliometric software has high standards and specifications for the data. Therefore, to ensure the quality and integrity of the collected data, we only searched the WoSCC database. The lack of a complement from other literature databases (such as Scopus), may have led to an incomplete analysis of the data. Secondly, this study only analyzed basic publication information and lacked in-depth exploration of the specific content, which is somewhat subjective. Finally, we only selected articles published in English; thus, significant articles published in other languages may have been excluded.

## Conclusions

This study analyzed 15 years of immune-related research on pancreatic cancer using VOSviewer and Citespace software, and systematically reviewed the trends in the field. The collaborative community of authors in this field is still in the process of formation, but there are already several well-known authors. The core journals that publish papers in this field are Cancers and Clinical Cancer Research. Scholars in the US publish the most articles, accounting for 44% of the articles, and are highly recognized in the field, in terms of the average number of citations per article. Keyword co-occurrence and cluster analyses revealed that several stable research themes have developed in this research field, such as the clinical treatment of pancreatic cancer, adoptive immunotherapy, etc. The author co-citation analysis revealed that the research hotspots in this field are constantly changing, and a careful analysis of the literature of highly cited authors may further explain the changing trend in hotspots.

## Supplemental Information

### Funding statement

This study is supported by National Natural Science Foundation of China (81573089,81972847); Tianjin key Medical Discipline (Specialty) Construction Project (TJYXZDXK-053B) and Tianjin Science and Technology Planning Project (21JCQNJC01900). The funders had no role in study design, data collection and analysis, decision to publish, or preparation of the manuscript.

### Additional Information and Declarations

All data presented in this study are included in this article and its supplementary files.

### Competing interests

The authors declare there are no competing interests.

### Author contributions

QX and YZ designed the study. QX, YZ and HZ collected the data. HZ, QX and YZ analyzed the data and drafted the manuscript. QX and YZ revised and approved the final version of the manuscript.

### Data availability

All data presented in this study are included in this article and its supplementary files.

## Notes

### Competing Interest Statement

The authors have declared no competing interest.

